# A genome-wide CRISPR screen in *Anopheles* mosquito cells identifies essential genes and required components of clodronate liposome function

**DOI:** 10.1101/2024.09.24.614595

**Authors:** Enzo Mameli, George-Rafael Samantsidis, Raghuvir Viswanatha, Hyeogsun Kwon, David R. Hall, Matthew Butnaru, Yanhui Hu, Stephanie E. Mohr, Norbert Perrimon, Ryan C. Smith

**Author notes:** These authors contributed equally and are listed alphabetically (Enzo Mameli, George-Rafael Samantsidis).

## Abstract

*Anopheles* mosquitoes are the sole vector of human malaria, the most burdensome vector-borne disease worldwide. Strategies aimed at reducing mosquito populations and limiting their ability to transmit disease show the most promise for disease control. Therefore, gaining an improved understanding of mosquito biology, and specifically that of the immune response, can aid efforts to develop new approaches that limit malaria transmission. Here, we use a genome-wide CRISPR screening approach for the first time in mosquito cells to identify essential genes in *Anopheles* and identify genes for which knockout confers resistance to clodronate liposomes, which have been widely used in mammals and arthropods to ablate immune cells. In the essential gene screen, we identified a set of 1280 *Anopheles* genes that are highly enriched for genes involved in fundamental cell processes. For the clodronate liposome screen, we identified several candidate resistance factors and confirm their roles in the uptake and processing of clodronate liposomes through *in vivo* validation in *Anopheles gambiae*, providing new mechanistic detail of phagolysosome formation and clodronate liposome function. In summary, we demonstrate the application of a genome-wide CRISPR knockout platform in a major malaria vector and the identification of genes that are important for fitness and immune-related processes.

## Introduction

Mosquitoes are essential vectors for the transmission of a variety of bacterial, viral, and parasitic pathogens that cause significant socioeconomic burden and mortality across the globe^1–3^. Among mosquito-borne diseases, malaria causes more than 200 million clinical cases and 600,000 deaths every year^4^, and is transmitted exclusively through the bite of an *Anopheles* mosquito. As a result of their public health importance, mosquitoes have become an emerging model system to examine aspects of development^5^, blood-feeding physiology^6^, vector-pathogen interactions^7^, and gene-drive technologies^8^; each with the ultimate goal of developing approaches to reduce the devastating impacts of mosquito-borne disease transmission.

There has been significant progress in our biological understanding of mosquito species through the development of genetic tools utilizing RNAi^9^, transgenesis^10,11^, and site-directed mutagenesis^12–15^. However, these reverse-genetic approaches only enable the investigation of candidate gene phenotypes. In contrast, the development of forward-genetic screen technologies would make it possible to associate genes with phenotypes in an unbiased manner and thereby uncover mosquito-specific as well as conserved gene functions. CRISPR gene editing technology has made it easier to perform genetics in mosquitos and other non-model species^16^, and CRISPR technologies are being applied as an *in vivo* research tool and potential intervention in mosquito populations, including mosquitoes of the *Anopheles* genus^17,18^. What has remained lacking, however, is an efficient system for genome-wide forward genetic screening using CRISPR or other similar technologies.

To address this need, we recently developed a platform for pooled-format CRISPR screening in mosquito cells, based on the CRISPR screen platforms we developed for *Drosophila* S2R+ cells^19,20^. For this approach, we use recombination mediated cassette exchange (RMCE) to integrate single guide RNAs (sgRNAs) into the genome, making it possible to later associate screen assay phenotypes with genotypes. Application of this approach in *Drosophila* cells has resulted in the identification of essential genes^19,21^, a novel transporter for the insect hormone ecdysone^22^, and receptors of bacterial toxins^23^. To extend this approach to *Anopheles*, we first engineered the *Anopheles* Sua-5b cell line with attP sites for RMCE and stable expression of Cas9 (i.e., a ‘screen-ready’ cell line); identified pol III promoters for sgRNA expression in *Anopheles* cells; and developed an approach to sgRNA design for screens in *Anopheles* Sua-5b cells. Then, in a pilot study, we introduced into the screen-ready Sua-5b cells a library of 3,487 sgRNAs and screened for cells resistant to treatment with rapamycin, ecdysone, or trametinib^24^. As expected, we were able to precisely and efficiently identify the *Anopheles* orthologs of the targets of these treatments^24^, opening the doors for the first time to application of large-scale forward-genetic screening in *Anopheles* cells.

One of our goals in developing the genome-wide cell screening platform was to contribute to our understanding of mosquito immune responses and cellular immune function^25^. Mosquito immune cells, known as hemocytes, are essential components of the innate immune system^26^ and have integral roles in shaping mosquito vector competence to both arbovirus^27,28^ and malaria parasite infection^29–33^. With few genetic resources available for the *in vivo* study of mosquito hemocytes, we recently adapted the use of clodronate liposomes, which have traditionally been used in mammalian systems for macrophage depletion^34,35^, to chemically ablate macrophage-like immune cell populations across arthropod species^32,36,37^. This methodology has been instrumental to our growing understanding of the role of macrophage-like granulocyte populations in mosquitoes and their contributions to host survival and pathogen infection outcomes^28,32,36^. However, despite the widespread use of clodronate liposomes in vertebrate and invertebrate systems, we still lack a mechanistic understanding of how they gain entry and are processed to promote targeted cell ablation. We reasoned that the application of a genome-wide CRISPR screen has the potential to identify factors that are required for clodronate liposome-mediated cell ablation, providing a methodology to better understand clodronate liposomes as a research tool.

Herein, we perform genome-wide CRISPR screens in an *Anopheles* mosquito cell line that identify ∼1300 essential genes responsible for cell viability and growth, as well as discern several genes involved in the uptake and processing of clodronate liposomes that provide novel insights into the mechanism by which they promote cell ablation. These results demonstrate the potential of forward-genetic screens in mosquito cell lines that have important implications for advancing our understanding of cellular immune function and the development of new mosquito control strategies.

## Results

### Genome-wide CRISPR knockout screen to identify essential genes in *Anopheles*

To extend the pooled screen approach to genome-wide scale, we first cloned a library of 89,711 unique sgRNAs targeting 93% of *Anopheles* genes, with ∼96% of these genes targeted by 7 sgRNAs per gene based on our previously reported sgRNA design resource for this species^24^ (**Fig. 1a** and **1b**). This set was supplemented with positive and negative control sgRNAs and others, resulting in a total library of 90,208 sgRNAs (**Supplementary Table 1**). We then introduced the library into CRISPR ‘screen-ready’ (attP+, Cas9+) *Anopheles* Sua-5b cells^24^ in the presence of ΦC31 integrase to generate a pool of knockout (KO) cells (**Fig. 1a**). Our first goal for genome-wide screening was to use a ‘dropout’ assay (negative selection assay) to identify genes for which knockout results in decreased fitness, growth arrest, and/or cell death (hereafter, “essential genes”). After 8 weeks of outgrowth of the KO cell pool, we compared the relative abundance of each sgRNA in the outgrowth pool to the distribution of sgRNAs in the starting plasmid library by NGS followed by MAGeCK MLE analysis^38^ (**Supplementary Table 2)**. Using the relationship between gene expression and Z-score rank, we identified 1280 putative essential genes with 95% confidence (FDR=0.05) (**Fig. 1c**). As expected, the majority of guides targeting genes annotated as components of the cytoplasmic or mitochondrial ribosome, the spliceosome, or the proteasome have negative Z-scores, consistent with essentiality in this assay (**Fig. 1d**).

**Fig. 1.**
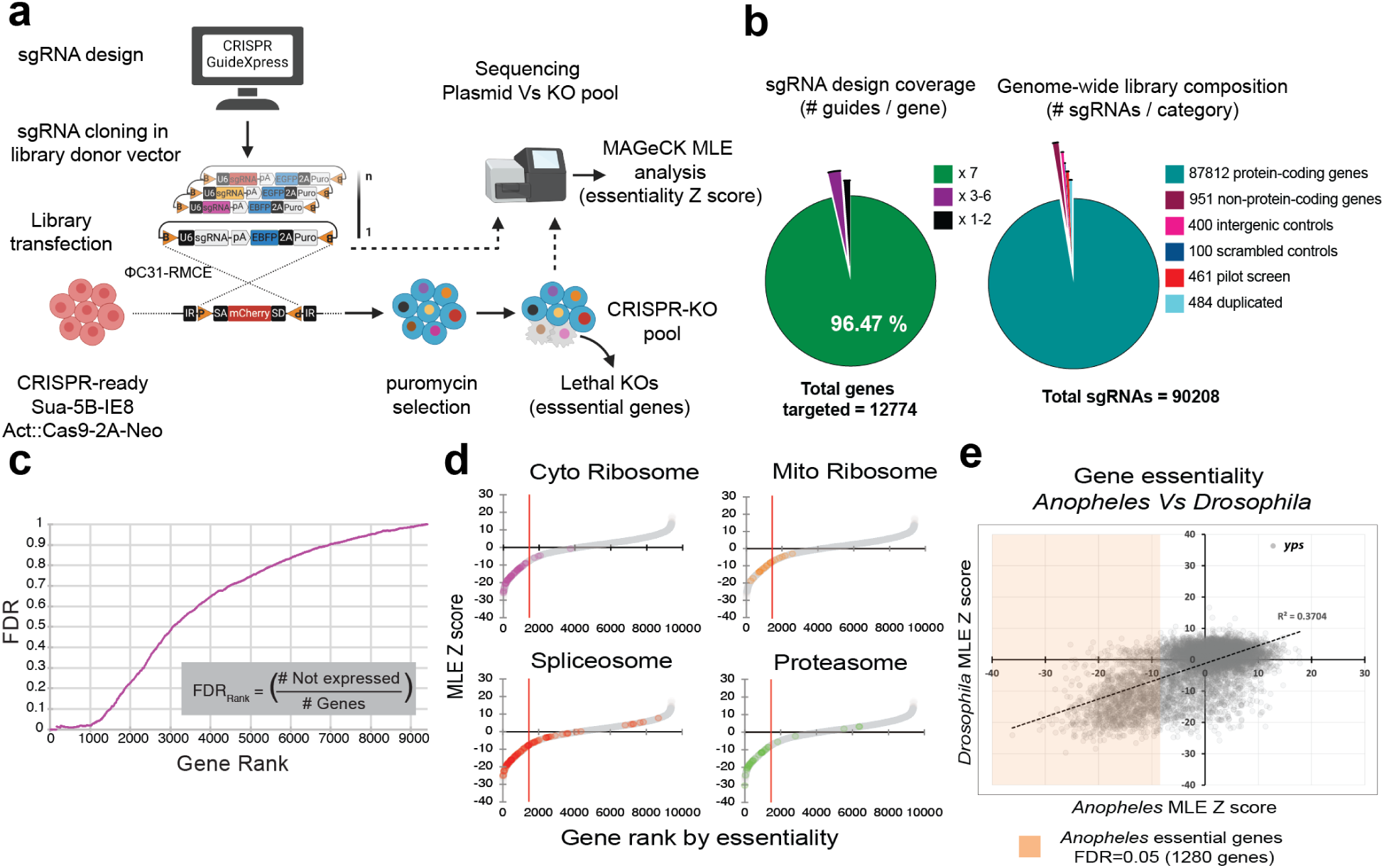
Genome-wide CRISPR knockout screen reveals genes required for fitness in *Anopheles* Sua-5B cells. (**a**) Schematic for gene essentiality CRISPR screen. CRISPR GuideXpress was used to design a whole-genome sgRNA library targeting protein-coding and non-coding *Anopheles gambiae* genes. The library was cloned into the pLib6.4B_695 vector and delivered to Sua-5B-IE8-Act::Cas9-2A-Neo via ΦC31 recombination-mediated cassette exchange to yield a pool of knockout cells. During outgrowth, cells that received sgRNAs targeting essential genes will “drop out” of the KO pool. The relative abundance of each sgRNA in the KO outgrowth pool of cells was compared to the plasmid library by NGS followed by MAGeCK MLE analysis. (**b**) Genome-wide library coverage. The starting library design includes 90,208 sgRNAs (88,763 unique sgRNAs) targeting 93% of *Anopheles* genes, with 7 sgRNAs per gene coverage for ∼96% of these genes. (**c**) Data analysis. MAGeCK MLE was used to analyze gene essentiality. Using the relationship between gene expression and Z-score rank, we can identify ∼1300 essential genes with 95% confidence (FDR=0.05). Their distribution by category is shown in (**d**): colored data points within the Z-score whole genome distribution (grey dots) highlight genes belonging to each essential category by Gene Ontology term (Cytoplasmic Ribosome KEGG:aga030008; Mitochondrial Ribosome GO:0098798,0005763; Spliceosome KEGG:aga03040; Proteasome KEGG:aga03050); red line intercept of *x axis* represents Z-score essentiality threshold at FDR=0.05. (**e**) Comparison of essential genes identified in *Anopheles* and *Drosophila*. To compare gene lists between the two species, all *Anopheles* genes were mapped to corresponding *Drosophila* orthologs, then their respective essentiality scores from cell pooled CRISPR knockout screens were plotted. Colored inbox within the plot highlights *Anopheles* essential genes with Z-scores within FDR=0.05; fitted linear trendline and R^2^ squared value are displayed to highlight the correlation trend between datasets; *yps* single datapoint was darkened to enhance its visibility.

To further examine the essential gene data set, we first identified *Drosophila* orthologs of the *Anopheles* genes identified in the screen, then performed gene set enrichment analysis (GSEA) using PANGEA^39^, a gene set enrichment tool that includes query of manually curated annotations for *Drosophila* (**Supplementary Fig. 1 and Supplementary Table 2**). We performed GSEA using generic gene ontology (GO) terms for biological process^40^ and as expected, found that the gene list is enriched for fundamental cellular processes such as DNA and RNA metabolism and cell cycle components (**Supplementary Fig. 1**). Next, we analyzed the list using as a reference the curated Gene List Annotation for *Drosophila* (GLAD) resource^41^ and similarly identified gene groups corresponding to fundamental activities or structures, e.g. components of the ribosome, proteasome, and spliceosome (**Supplementary Fig. 1**). Finally, we used PANGEA to perform GSEA using as a reference a set of phenotypes associated with classical mutations as annotated by FlyBase^42^. Consistent with our expectations, the top-enriched phenotype is “cell lethal.” In addition, we found other enriched cell phenotypes, including “decreased occurrence of cell division,” “abnormal cell cycle,” and “abnormal cell size” (**Supplementary Fig. 1**).

Strikingly, we identified a single gene, *ypsilon schachtel* (*yps*) (AGAP006108; FBgn0222959), a ribonucleoprotein complex component^43^, that appears to limit cell growth in both *Anopheles* and *Drosophila* cells^19^, as suggested by the notable growth advantage in *yps* knockout cells (**Fig. 1e**, upper-right quadrant). Included among the *Anopheles* genes that negatively impacted growth, we identified the ortholog of *Drosophila serpent* (*srp*; AGAP002238; FBgn0003507), a GATA transcription factor involved in *Drosophila* hematopoiesis^44,45^. When *srp* was silenced *in vivo* in adult female *Anopheles gambiae*, we see reduced hemocyte numbers and increased malaria parasite infection (**Supplementary Fig. 2**), supporting that srp has similar roles in mosquito hematopoiesis and immune function.

### Comparison with essential gene screen data from *Drosophila* and human cells

As a data quality analysis step, we next compared putative essential genes identified in this screen with essential genes identified in a similar screen in *Drosophila*^21^. To do this, we first mapped *Anopheles* genes to *Drosophila* orthologs using DIOPT (v 9.0) and filtered the results based on the DIOPT score. For genes in each ortholog pair, we graphed the corresponding Z scores and found a high degree of overlap between genes that scored as essential in the two species (**Fig. 1e**, lower-left quadrant), supporting the validity of the results of the essential gene screening platform in *Anopheles* cells. We next used the list of *Drosophila* orthologs to ask how many genes are in common in the *Anopheles* essential gene list and a similar list generated using an optimized CRISPR knockout screen platform in *Drosophila* S2R+^21^. The 1280 mosquito genes map to 1213 *Drosophila* genes and of these, 88% (1073/1213) were identified as essential in the *Drosophila* cell screen (**Supplementary Table 3**).

The results of comparison of essential genes in *Anopheles* and *Drosophila* cell screen datasets suggests that many of the genes are generally required for cell growth and viability but is confounded by the fact that both *Anopheles* Sua-5b and *Drosophila* S2R+ cell lines are considered hemocyte-like (blood-like) cell types, such that conserved factors essential for insect cell hemocytes could be included in both lists. To explore this further, we next mapped genes on the *Anopheles* essential gene list to human orthologs, and asked how many of these genes are included in a core list of 684 human cell-essential genes compiled based on data from 17 human cell knockout screens^46^. The 1280 *Anopheles* essential genes mapped to 1185 human orthologs and of these, 34% (398/1185) are among the core human essential genes (**Supplementary Table 3**), suggesting that these 398 genes are conserved genes essential in distantly related metazoan cells.

### Genome-wide CRISPR screen for resistance to clodronate treatment

Recent studies have demonstrated the use of clodronate liposomes as a valuable tool to probe cellular immune function in arthropods^32,36,37^, yet at present, we lack a fundamental understanding of how they function. Even in vertebrate systems, where clodronate liposomes have been more widely used^34,35^, there is only limited mechanistic information as to how these particles function^47^. As a result, we reasoned that screening for resistance to clodronate liposome-mediated cell ablation in *Anopheles* Sua-5b cells, a hemocyte-like cell line^48^, could reveal important factors relevant to clodronate liposome function in mosquito immune cells. To initiate a genome-wide clodronate liposome selection-based screen, we first tested the effects of treatment of screen-ready Sua-5b cells with a range of concentrations of clodronate liposomes or control (empty) liposomes to determine the appropriate concentrations for a selection-based screen (**Fig. 2a**). We found that the IC50 of the clodronate liposomes for Sua-5b cells was 7.4 μM, whereas the IC50 of control liposomes was approximately 11-fold higher (81.6 μM; **Fig. 2a**). To perform the screen, we subjected a pooled library of Sua-5b KO cells to continuous selection with clodronate liposomes (“Clodronate A” group), treated them for 4 days with clodronate liposomes then followed by outgrowth in standard media (“Clodronate B” group), or treated them continuously with control liposomes for a total of three cycles of treatment/outgrowth (**Fig. 2b**). Following the last cycle of outgrowth, we used deep amplicon sequencing and MAGeCK analysis^38^ to compare sgRNA abundance in each of the two experimental and the control population **(Supplementary Table 4)**.

**Fig. 2.**
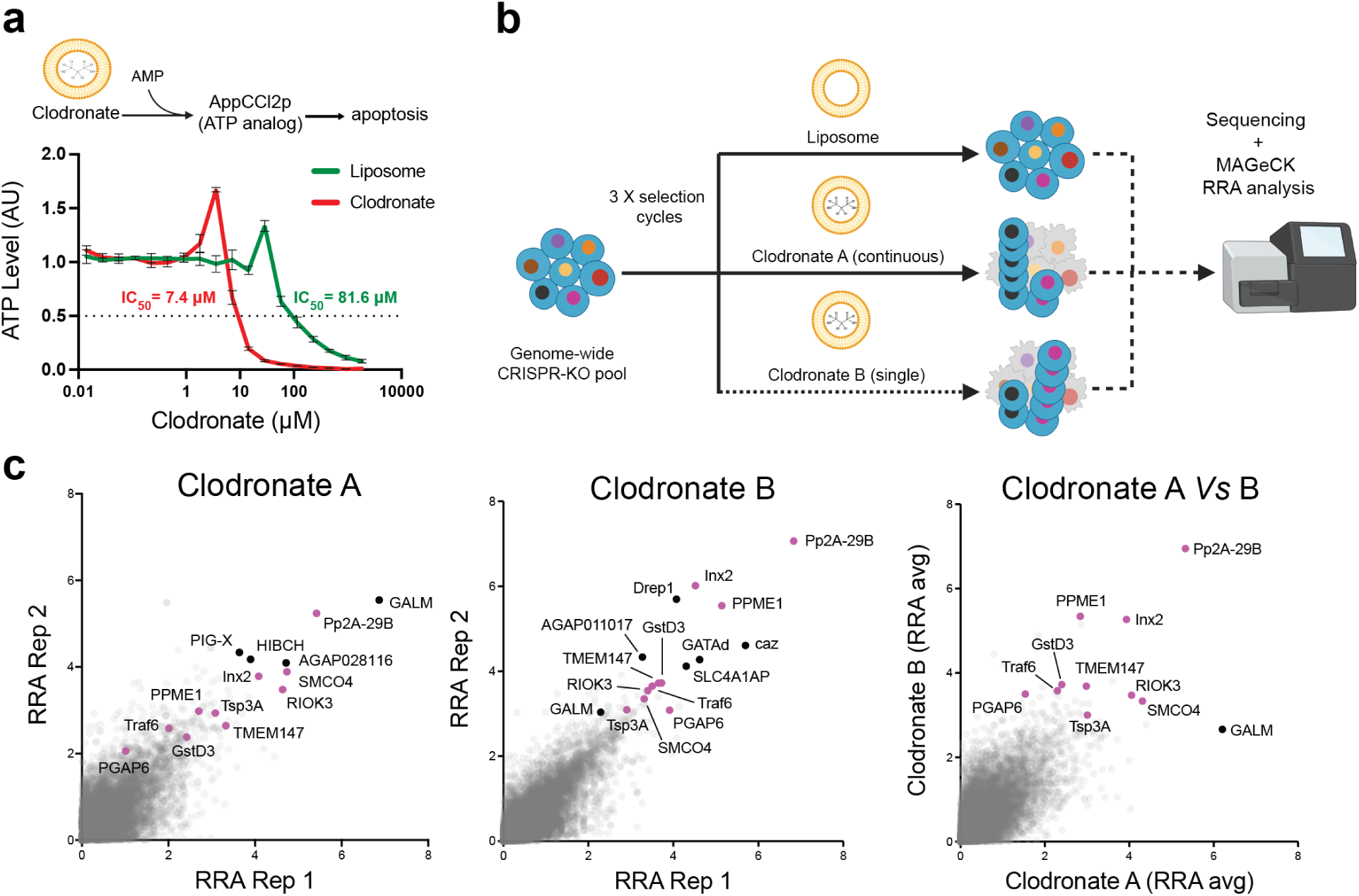
Genome-wide CRISPR knockout screen reveals genes for which knockout confers resistance to clodronate liposome uptake and/or induced cell death in *Anopheles* cells. **(a)** Clodronate liposomes induce cell death after cellular uptake by releasing clodronate, which is enzymatically converted to adenosine 5’β-γ-dichloromethylene triphosphate (AppCCl2p), an ATP analog that can induce apoptosis. Assay of total ATP levels reveals higher lethality in the *Anopheles* Sua-5B-IE8-Act::Cas9-2A-Neo cells treated with clodronate as compared to a liposome control (11-fold difference in relative IC50 values). (**b**) Schematic of a genome-wide positive selection CRISPR knockout screen for clodronate resistance. A genome-wide CRISPR KO pool of Sua-5B-IE8-Act::Cas9-2A-Neo cells was left untreated or treated with 16 μM liposome (control) or 8μM clodronate. All treatments were performed for three cycles and with continuous drug selection of the KO pool, except for the “clodronate B” treatment group, for which cells were subjected to an initial treatment for 4 days and then allowed to recover in non-selective media at each treatment cycle. Genomic DNA from endpoint cell populations was used for PCR amplification of sgRNAs, followed by NGS and enrichment analysis using the MAGeCK robust rank aggregation (RRA) algorithm. (**c**) Scatter plots of RRA scores for two replicates, comparing clodronate treatments A and B to control liposome treatments (left and center panels) or comparing average RRA scores between the two treatments (right panel). Left and center panels: The ‘hits’ (positive results) chosen for follow-up studies are labeled with gene symbols. Hits shown in black represent the top eight genes by RRA rank. Hits in magenta are genes of interest selected from among the top 50 hits from each screen. Right panel: only genes of interest and top hit gene of the clodronate A screen (in black) are indicated. Gene symbols shown are the symbols for orthologous human genes (symbols in all caps) or orthologous *Drosophila* genes.

To identify candidate genes involved in clodronate uptake and/or processing, we compared the liposome control to the clodronate treatment groups (i.e., we compared Clodronate A or Clodronate B treatments to the liposome control). We were able to identify genes enriched in the experimental groups (**Fig. 2c**). The top-scoring gene in the continuous treatment (Clodronate A) group is a predicted *Anopheles* ortholog of the mammalian GALM (AGAP008154), whereas the top-scoring gene in the Clodronate B group is a predicted ortholog of mammalian PPP2R1A and Pp2A-29B in *Drosophila* (herein referred to as Pp2A-29B; AGAP009105). While some top-scoring genes were different between the two treatment groups, twelve genes scoring in the top 50 hits were found in common between both screens (**Fig. 2c**). This includes Pp2a-29B, PPME1 (AGAP008336), SMCO4 (AGAP003534), RIOK3 (AGAP009993), Inx2 (AGAP001488), PAFAH1B2 (AGAP000939), Tsp3A (AGAP002257), TMEM147 (AGAP008757), FAM117B (AGAP011572), jbug (AGAP007006), caz (AGAP001645), and AGAP011017. To reveal the genetic determinants of clodronate liposome uptake and induced toxicity, we performed similar GSEA analyses as for the essential gene set (**Supplementary Table 2**), on the top scoring genes conferring resistance to clodronate liposome treatment from each screen. GSEA was performed with GO biological process terms from standard GO sets (“GO hierarchy” at PANGEA); GO subsets specifically curated for *Drosophila* by Flybase^42^ and the Alliance for Genome Resources^49^ (Slim2); and FlyBase Gene Groups^50^. A common theme that emerged from our GSEAS analysis was the enrichment for methyltransferases and gene sets enriched in the *Drosophila* GO analysis included an “autophagy” gene set (**Fig. S3** and **Supplementary Table 4**).

### Optimization of clodronate liposome concentrations and timing of uptake *in vivo*

Previous *in vivo* studies using clodronate liposomes in *An. gambiae* were performed using a concentration of ∼120 μM/ml (1:5 dilution)^32,33,51^, a concentration much higher than the ∼8 μM/ml concentration used herein for our *in vitro* screening experiments (**Fig. 2a**). To confirm that this lower concentration was still able to promote cell ablation *in vivo*, we compared the efficiency of clodronate liposomes at the 1:5 dilution with that of a 1:50 dilution (∼12 μM/ml; comparable to that used in *in vitro* experiments). Using the expression of *eater* and *Nimrod B2* as a proxy for mosquito immune cell (granulocyte) numbers as previously^32,33,36,37,51^, both the 1:5 and 1:50 clodronate liposome dilutions were able to promote similar reductions in *eater* and *Nimrod B2* (**Supplementary Fig. 4**), suggesting that both concentrations were equally effective in their ability to reduce mosquito immune cell populations *in vivo*.

Similarly, while previous studies have demonstrated the utility of clodronate liposomes to deplete immune cell populations in flies, mosquitoes, and ticks^32,33,36,37,51^, the precise timing required for phagocyte depletion has not been previously examined. Therefore, we utilized fluorescent liposome particles (LP-DiO) to determine the temporal kinetics of liposome uptake and subsequent phagocyte depletion. When examined at multiple time points after injection, the uptake of fluorescent LP-DiO particles peaked at 6h post-injection (with ∼37% of hemocytes LP-DiO^+^), before the percentage of LP-DiO^+^ cells began to decrease over time (**Supplementary Fig. 5**). To validate these findings in the context of clodronate-mediated phagocyte depletion, we performed similar time-course experiments following the injection of control or clodronate liposomes to evaluate the timing needed to initiate phagocyte depletion. When granulocyte numbers were assessed by proxy via qPCR through the expression of *eater* and *Nimrod B2*^32,33,36,37,51^, there was no effect on granulocyte numbers at 6 hours post-injection, yet by 8 hours there was a significant and sustained reduction in *eater* and *Nimrod B2* transcripts indicative of granulocyte depletion (**Supplementary Fig. 5**). Together, these data suggest that liposome uptake occurs within hours post-injection and that liposomes are quickly processed to promote phagocyte depletion. Since previous studies have only evaluated phagocyte depletion at 24 or 48h post-injection^32,33,36,37,51^, these data provide greater resolution into the timing of liposome processing, enabling a more precise evaluation of candidate genes identified in our CRISPR screen to examine clodronate liposome function.

### *In vivo* validation of candidate genes

We next identified candidates from the CRISPR cell screen (**Fig. 2**) for further validation *in vivo* in *An. gambiae* hemocytes. To do this, the top 50 hits from each replicate (of which 12 genes were identified in both screens) were cross-referenced with a previous scRNA-seq of *An. gambiae* hemocytes^51^ to confirm their expression in mosquito granulocyte populations (**Fig. 3a**). Candidates were selected for further analysis based on their presence in both screens and predicted functional annotations (**Fig. 2c**, **Supplementary Table 4**). A total of 10 candidates were selected for further validation *in vivo* (**Fig. 3a**) using RNA interference (RNAi). To evaluate the role of each candidate gene, we performed dsRNA injections for all 10 candidate genes, resulting in the successful knockdown of 5 out of the 10 genes (*Tsp3A*; *PGAP6,* AGAP002672; *Traf6*, AGAP003004; *GstD3*, AGAP004382; *TMEM147*) when evaluated at two days post-injection (**Fig. 3b**). Additional experiments to examine gene-silencing at four days post-injection for the remaining candidates similarly failed to induce a knockdown (**Supplementary Fig. 6**), suggesting that these genes are not amenable to gene-silencing.

**Fig. 3.**
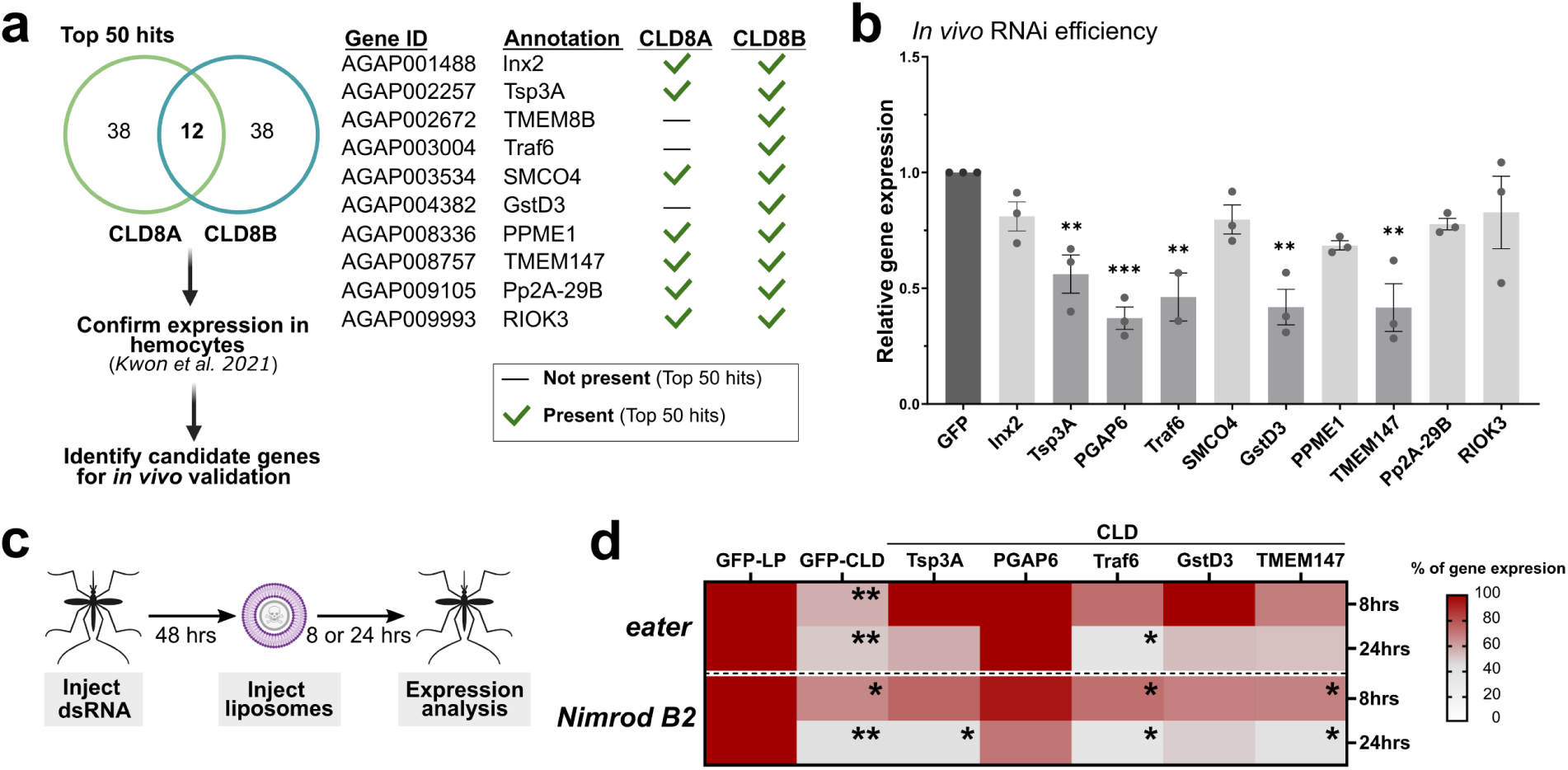
RNAi and *in vivo* validation of candidate genes involved in clodronate liposome function. (**a**) Genes identified from both clodronate liposome screens were selected based on their enrichment in both screens, expression in mosquito hemocyte populations^51^, and presumed biological function to select for candidate genes for further validation *in vivo*. Following the injection of gene-specific dsRNAs, the efficiency of RNAi was evaluated in whole mosquitoes via qRT-PCR two days post-injection (**b**). Expression data from three independent experiments are displayed as the mean ±SEM and compared to GFP controls. Statistical differences were examined using an unpaired t-test for each individual gene compared to controls. (**c**) To determine the influence of candidate genes on clodronate liposome function, candidate genes were first silenced via the injection of dsRNAs, then control or clodronate liposomes were injected two days post-dsRNA injection. The effects of gene-silencing on clodronate liposome function were assessed by the expression of *eater* and *Nimrod B2* as a proxy to measure immune cell depletion (**d**). The heatmap summarizes the effects of gene-silencing on the efficacy of clodronate liposome-mediated cell ablation at 8 and 24 hours, where non-significant changes in *eater* and *Nimrod B2* expression support that the gene-silenced background impairs clodronate liposome function. Data represent three or more independent experiments. For each RNAi background, expression data were compared between control liposomes and clodronate liposomes. Differences in gene expression were examined using an unpaired t-test. For all experiments, significant differences are indicated by asterisks (*, *P* < 0.05; **, *P* < 0.01; ***, *P* < 0.001).

To confirm candidate gene function in clodronate liposome-mediated phagocyte depletion, RNAi was performed in adult female mosquitoes before injection with control or clodronate liposomes. The influence of RNAi on clodronate-mediated granulocyte depletion was then evaluated at 8 or 24 hours via the expression of *eater* and *Nimrod B2* as a proxy of granulocyte numbers^32,33,36,37,51^ (**Fig. 3c**). While clodronate liposome treatment significantly reduced *eater* and *Nimrod B2* expression at both 8- and 24-hours post-injection in dsGFP controls (**Fig. 3d**), silencing of *Tsp3A*, *PGAP6*, *Traf6*, *GstD3*, and *TMEM147* each impaired phagocyte depletion, resulting in higher expression levels of *eater* and *Nimrod B2* when compared to controls (**Fig. 3d**). There was variance amongst the five candidate genes examined in their effects on phagocyte depletion, with the silencing of *Traf6* displaying the weakest phenotype (only affecting *eater* at 8 hours), while *PGAP6* silencing completely inhibited the effects of clodronate liposome treatment at 8 and 24 hours for both reporter genes examined (**Fig. 3d**). Together, these phenotypes confirm the role of each candidate gene in clodronate liposome-mediated phagocyte depletion.

### Liposome uptake is mediated by phagocytosis

To better understand the roles of our candidate genes and the uptake mechanisms of clodronate liposomes in invertebrate cells, we first examined the influence of endocytic pathways on liposome uptake. Using pharmacological inhibitors that target endocytosis (chlorpromazine, CPZ)^52–54^ or phagocytosis (cytochalasin, CytoD)^55–58^ (**Fig. 4a**), mosquitoes were intrathoracically injected with each inhibitor or 10% DMSO as a control to determine the role of each respective pathway on liposome uptake. When mosquitoes were challenged with LP-DiO particles following inhibitor treatment, the uptake of LP-DiO particles was significantly impaired only in mosquitoes treated with CytoD (**Fig. 4b**), suggesting that liposome uptake is dependent on immune cell phagocytosis. Additional experiments confirm that CytoD treatment impairs phagocyte depletion (**Fig. 4c**), demonstrating that phagocytic function is integral to clodronate liposome-mediated phagocyte depletion.

**Fig. 4.**
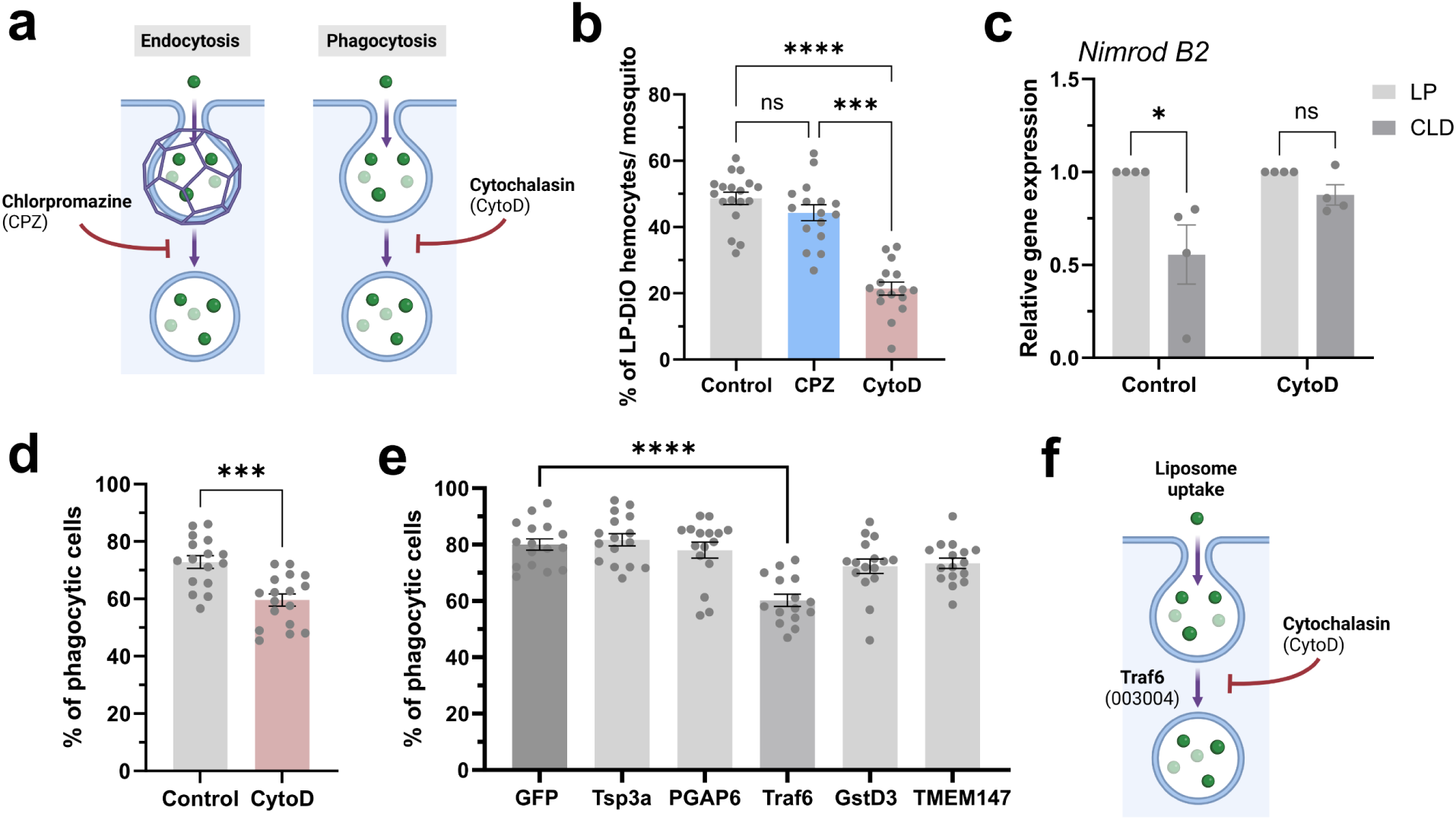
Clodronate liposome uptake is mediated by phagocytosis. (**a**) Overview of endocytic pathways, clathrin-mediated endocytosis and phagocytosis, with their respective inhibitors. To address the manner by which lipsomes undergo uptake, mosquitoes were injected with the respective endocytic inhibitors (**a**), with the uptake of LP-DiO particles by mosquito hemocytes (immune cells) assessed at 8 hours post-injection as the percentage of LP-DiO^+^ hemocytes (**b**). Data were collected from individual mosquitoes (dots) and examined using Kruskal-Wallis with a Dunn’s multiple comparisons test. (**c**) The effects of phagocytosis-inhibition (via cytochalasin D, cytoD) on clodoronate liposome efficacy were evaluated using *Nimrod B2* expression as a proxy for immune cell ablation. Data from four independent experiments were analyzed using multiple unpaired t-tests to determine significance. After confirmation that cytoD treatment reduces the uptake of fluorospheres (**d**), candidate genes from the CRISPR screen were evaluated for potential phenotypes that similarly influence phagocytosis (**e**). Data from **d** and **e** display values collected from individual mosquitoes, with analysis respectively performed with individual (Mann-Whitney) or multiple comparisons (Kruskal-Wallis andDunn’s multiple comparisons test) to determine significance. These data contribute to a model (**f**) suggesting that liposome uptake is mediated by phagocytosis and implicates Traf6 (with AGAP gene ID number in parenthesis) in phagocytic uptake. For all experiments, significant differences are indicated by asterisks (*, *P* < 0.05; ***, *P* < 0.001; ****, *P* < 0.0001). Summary figures created with BioRender.com.

After demonstrating that CytoD treatment impedes phagocytosis *in vivo* (**Fig. 4d**), we sought to address whether any candidate genes identified in the CRISPR screen may similarly influence phagocytosis and liposome uptake. When phagocytosis experiments were performed following RNAi-mediated gene silencing, only the *Traf6*-silenced background displayed notable defects in phagocytic ability (**Fig. 4e**). This suggests that the impairment of clodronate liposome-mediated phagocyte depletion by *Traf6* RNAi (**Figs. 3a** and **3d, Supplementary Table 4**) is likely mediated through phagocytic function (**Fig. 4f**). Moreover, the minimal influence of the remaining candidate genes on phagocytosis suggests that their function lies downstream of liposome uptake.

### Candidate genes that impair clodronate liposome processing are involved in phagolysosome formation

Following phagocytic uptake, internalization results in the formation of an early phagosome that undergoes maturation and ultimately fuses with the lysosome to form a phagolysosome, facilitating pathogen killing and protein degradation^59–61^ (**Fig. 5a**). To better understand how clodronate liposomes are processed following phagocytosis and to identify potential roles of our candidate genes in this process, we again utilized LP-DiO particles to visualize liposome uptake and processing in mosquito immune cells. Approximately 8 hours post-injection, LP-DiO particles colocalize with lysosomes (**Fig. 5b**), indicating that the normal processing of liposome particles involves the formation of the phagolysosome (**Fig. 5a**). In addition, we observed distinct patterns of DiO localization in immune cells, with some cells displaying punctate DiO localization, suggesting the presence of intact LP-DiO particles (referred to as LP-DiO+ cells, **Fig. 5c**), or those that displayed a more diffuse pattern of DiO suggesting the breakdown and release of the LP-DiO particles (referred to as DiO+ cells, **Fig. 5d**). When these phenotypes were quantified in our candidate gene backgrounds, both *Tsp3A* and *Traf6* RNAi displayed a significant increase in the accumulation of LP-DiO+ cells (**Fig. 5c**). Conversely, silencing of *Tsp3A*, *PGAP6*, and *TMEM147* significantly reduced the percentage of cells that were DiO+ (**Fig. 5d**), suggesting that these RNAi backgrounds were impaired in their abilit to breakdown LP-DiO+ particles. Together, these data suggest that each of our candidate genes, with the exception of *GstD3*, contribute to the internal processing of liposome particles likely through the formation of the phagolysosome.

**Fig. 5.**
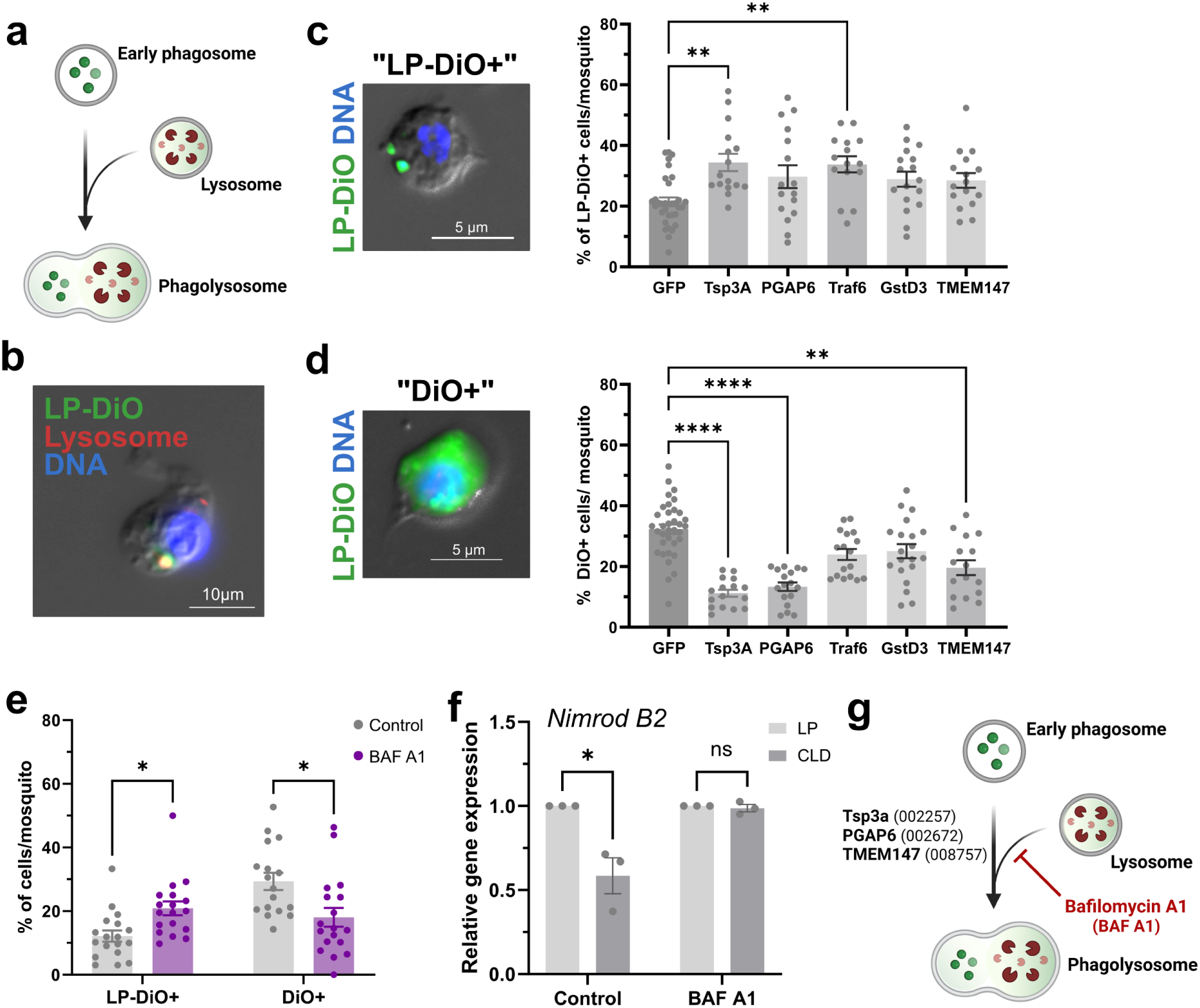
Clodronate liposome processing requires candidate genes involved in phagolysosome formation. (**a**) Overview of phagosome maturation and phagolysosome formation after fusion with the lysosome. To confirm that phagocytosed liposome particles undergo phagolysosome formation, immunofluorescence assays were performed following LP-DiO injection and staining with lysosome-specific dye, LysoView 594 (**b**). Co-localization of LP-DiO particles (green) and the lysosome (red) support that liposomes are processed by the formation of the phagolysosome prior to degradation. Observations of LP-DiO^+^ cells display two distinct phenotypes, where LP-DiO particles are punctate and remain intact (referred to as LP-DiO^+^; **c**), or where DiO fluorescence is diffused (referred to as DiO^+^) suggesting that liposome particles have been degraded (**d**). These LP-DiO^+^ (**c**) and DiO^+^ (**d**) phenotypes were evaluated in the gene-silenced backgrounds for each of the candidate genes identified in the CRISPR screen. Data were collected from individual mosquitoes (dots) and examined using Kruskal-Wallis with a Dunn’s multiple comparisons test. (**e**) To further refine these observed phenotypes, we evaluated LP-DiO^+^ and DiO^+^ phenotypes following treatment with Bafilomycin A1 (BAF A1), an inhibitor of lysosome fusion with the phagosome. Data are displayed from individual mosquitoes (dots). (**f**) The effects of BAF A1 inhibition on clodronate liposome function were evaluated from three independent experiments using *Nimrod B2* expression as a proxy for immune cell ablation. Data in **e** and **f** were analyzed using multiple unpaired t tests to determine significance. These data contribute to a model (**g**) suggesting that liposome processing is mediated by formation of the phagolysosome involving Tsp3A, PGAP6, and TMEM147, which can be impaired using the inhibitor BAF A1. For all experiments, significant differences are indicated by asterisks (*, *P* < 0.05; **, *P* < 0.01; ****, *P* < 0.0001). Summary figures created with BioRender.com.

To further validate this phenotype, we performed additional experiments using Bafilomycin A1 (BAF A1), an inhibitor of lysosome acidification and phagolysosome formation (**Fig. 5a**). Similar to the DiO localization phenotypes observed in **Figs. 5c** and **5d**, BAF A1 treatment significantly increased the percentage of LP-DiO+ cells, while reducing the percentage of DiO+ cells (**Fig. 5e**). Additional experiments to evaluate clodronate liposome function in the BAF A1-treated background demonstrated that BAF A1 significantly inhibits clodronate liposome-mediated phagocyte depletion (**Fig. 5f**). The observed phenotypes are strikingly similar to the *Tsp3A*-silenced background, as well as the partial phenotypes associated with *PGAP6, Traf6,* and *TMEM147* RNAi which support the hypothesis that these candidate genes have essential functions in phagolysosome formation (**Fig. 5g**).

Together, these data support a model in which the phagocytic uptake of liposomes involves Traf6 and can be inhibited by CytoD treatment (**Fig. 6**). Additionally, the knockdown of several genes, such as Tsp3a, PGAP6, and TMEM147, mimics the effect of the BAF A1 inhibitor, indicating their role in further liposome processing and phagolysosome formation.(**Fig. 6**). Although silencing of *GstD3* influenced clodronate liposome function (**Fig. 3**), experiments examining liposome uptake and processing did not yield phenotypes for *GstD3*, suggesting that GstD3 contributes to the downstream events that promote cell ablation (**Fig. 6**).

**Fig. 6.**
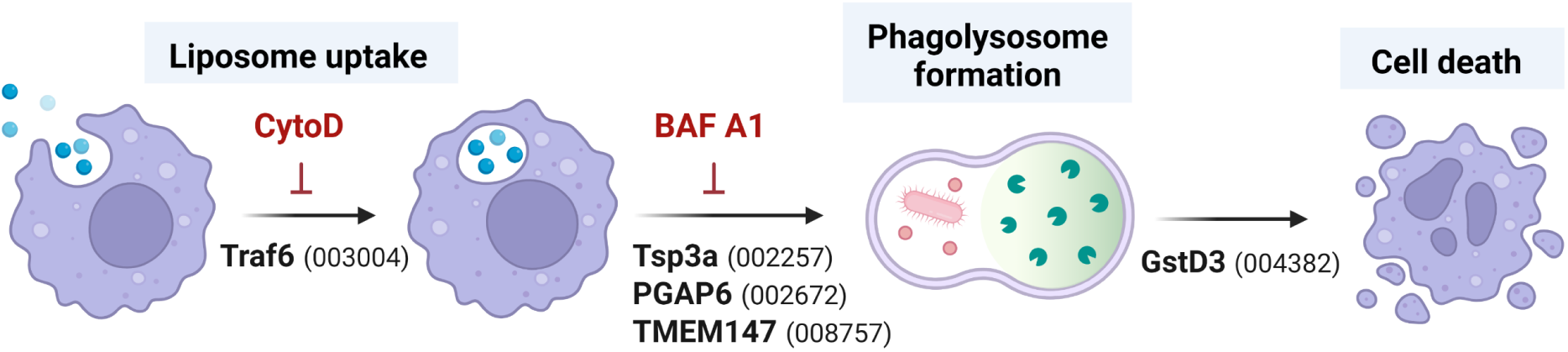
Summary of candidate genes involved in clodronate liposome function. Experiments support a model in which candidate genes identified in the CRISPR screen contribute to the uptake, processing, and downstream function of clodronate liposomes in promoting immune cell ablation in *An. gambiae*. Figure created with BioRender.com.

## Discussion

Forward-genetic CRISPR knockout screens enable an unbiased interrogation of gene function across a wide range of biological topics^62^. Although evidence has demonstrated the utility of this forward-genetics approach from mammals^63,64^ to *Drosophila*^19,22^, the methodology had yet to be fully extended to other insect systems. We previously developed the technology to enable pooled CRISPR knockout screening in mosquito species and demonstrated its application through initial proof-of-principle studies^24^. Here, we performed the first genome-wide pooled CRISPR screens in *Anopheles* to identify genes with essential roles in host fitness and provide new insights into the mechanisms of clodronate liposome function in mosquitoes.

Our genome-wide CRISPR knockout fitness screen identified a total of 1280 genes using a 5% FDR cutoff that are required for *Anopheles* Sua-5b cell growth, division, and/or viability. Most of these genes (88%) are also essential in *Drosophila* S2R+ cells^21^, and are highly enriched for genes encoding proteins involved in fundamental cell functions, such as protein synthesis, RNA splicing, and protein degradation (**Fig. 1d**). In addition, the list of *Drosophila* orthologs of the *Anopheles* essential genes includes 20% of genes annotated in the GLAD resource as “mitochondrial,” 18% of genes annotated as “metabolic,” 16% of genes annotated as “RNA-binding,” and 12% of genes annotated as “transcription factors” (**Supplementary Table 2**). Furthermore, we found significant overlap between the mosquito essential gene list and a list of ‘core essential genes’ identified in 17 CRISPR knockout screens in human cells^46^. Altogether, these findings support the quality of the gene dataset for mosquitoes and help to define a core set of essential genes shared across metazoa.

Notably, the essential *Anopheles* genes identified in this screen might help lead to the development of new approaches for mosquito control, such as targets for population suppression strategies that aim to reduce or eliminate mosquito populations^65^. For example, an essential *Anopheles* gene could be genetically targeted to create a synthetic gene-drive system capable of promoting lethality. In addition, the targeting of an essential mosquito gene has the potential to enhance population replacement strategies relying on CRISPR-Cas9^17^, homing endonuclease^66^, Medea-like^67,68^, or cleave and rescue^69,70^ as a means for selection against non-replacement alleles in split-drive systems.

Clodronate liposomes have been widely used in studies of vertebrate immunology^34,35^, and more recently in arthropod systems^32,36,37^, to promote the targeted ablation of phagocytic immune cell populations. While evidence suggests that clodronate-derived metabolites act as ATP analogs to block mitochondrial ATP synthase activity and consequently trigger apoptosis^47^, the precise mechanisms of clodronate liposome uptake and processing have not been adequately explored. Herein, the results of our genome-wide CRISPR screen provide a comprehensive examination of clodronate liposome function in *An. gambiae*. Using two screening methodologies, we identify a core set of 88 genes that are identified in one or both of our screens. This includes the enrichment of genes involved in cellular metabolism, methyltransferase function, and autophagy that bring new mechanistic insight into clodronate liposome function.

While the lack of RNAi and other limitations prevented downstream experiments for all hits identified in the clodronate liposome screen, a further examination of several identified genes infer additional subcellular components and pathways involved in clodronate liposome function. For example, multiple hits are components of or involved in the regulation of the serine/threonine-protein phosphatase 2A (PP2A) protein complex involved in a variety of biological processes such as cell growth, differentiation, apoptosis, and immune regulation^71^ PPME1, a methyl-esterase enzyme that acts directly on the catalytic subunit by demethylation of the PP2A protein complex to cause its inactivation^72^. Two other hits contribute to the regulation of the same protein complex: LCMT1 (AGAP008768), a methyl-transferase enzyme responsible for the methylation of PP2A at the same site targeted by PPME1^73^, and MASTL (AGAP001636) that acts by indirectly promoting the inactivation of PP2A^74^. With the PP2A protein phosphatase complex a master regulator of several cellular functions, it could be important for clodronate induced toxicity by mediating cytoskeleton rearrangements important for the uptake, trafficking, or degradation of liposomes, to the downstream steps controlling its toxicity by activating the Toll Like Receptor 3 (TLR3) cascade and apoptosis. Of note, both Pp2A-29B and MASTL are essential genes in human cells (DepMap) and in *Anopheles* cells, yet are among the most enriched targets in both clodronate screens. This suggests that, although the knockout of these genes impacts cell fitness under “normal” conditions, their knockout provides a growth advantage under clodronate selection, as the cells become less sensitive to the drug compared to normal cells. In addition, multiple hits that correspond to a serine/threonine-protein kinase signaling pathway involving RIOK3, ribosome biogenesis and regulation of type I interferon (IFN)-dependent immune response are represented in our clodronate liposome screen. In addition to RIOK3, two other hits possibly belong in the same pathway, Oseg4 (AGAP011562), a TNF-stimulated gene able to induce caspase 3-mediated apoptosis^75^ and RPS17, a component of the 40S ribosomal subunit that directly interacts with RIOK3 during ribosome biogenesis^76^. However, further experiments are required to establish exactly how these respective PP2A and RIOK3 signaling components are involved in clodronate liposome function.

Through the use of pharmacological inhibitors that target endocytic pathways, we demonstrate that the cellular uptake of clodronate liposomes is mediated by phagocytosis, and not clathrin-mediated endocytosis, providing further support for the specificity of clodronate liposomes to explicitly target phagocytic immune cells in both arthropods^36^ and mammals^77^. In addition, one candidate identified in our clodronate liposome CRISPR screen, Traf6, displayed notable defects in phagocytosis following *Traf6*-silencing, supporting that Traf6 likely influences the uptake of clodronate liposomes. However, as a RING-type ubiquitin ligase that interacts with several immune signaling molecules^78,79^, these effects are likely indirect. As a result, the phenotypes associated with *Traf6*-silencing may be caused by the impaired production of downstream immune effectors or defects in immune cell activation^80^.

Additional microscopy, RNAi, and inhibitor experiments confirm that the formation of the phagolysosome is a critical step in the processing of clodronate liposomes. Lysosomes contain various hydrolytic enzymes that promote the breakdown of macromolecules for degradation and cellular recycling^81^, thereby serving an essential role in the breakdown of the liposome particle and the intracellular delivery of clodronate required to initiate cell death. Taking advantage of the fluorescence of LP-DiO particles, we demonstrate the co-localization of liposome particles with the lysosome, as well as the punctate and diffuse patterns of DiO that enable the visualization of liposome processing. We demonstrate that three candidate genes identified in our CRISPR screen, *Tsp3A*, *PGAP6*, and *TMEM147*, have key roles in phagolysosome formation and validate these phenotypes in liposome degradation through the use of the BAF A1 inhibitor to impair lysosome fusion. While these data implicate Tsp3A, PGAP6, and TMEM147 in the intracellular processing of clodronate liposomes, their exact functions could not be fully resolved in our study. Each of these genes are believed to localize to cell membranes and have been implicated in immune cell function in orthologous systems^82–84^. While additional details of Tsp3a and TMEM147 function are limited, the human ortholog of PGAP6 is a GPI-anchored phospholipase with predicted localization to the lysosome^83,85^, suggesting that PGAP6 could be essential to the breakdown of liposome particles and the subsequent release of clodronate following phagolysosome formation. Similarly, other genes identified in our screen such as CLVS1 (AGAP005388) that are required for proper formation of late endosomes and lysosomes^86^, and AGAP011017 which is of unknown function and harbors a putative lipid binding domain (InterPro) similar to the Ganglioside GM2 activator (GM2-AP) that acts as a lysosomal lipid transfer protein, further implicate lysosome fusion as an important step in clodronate liposome function. However, one limitation of these experiments was our inability to further define the role of GstD3 in clodronate liposome function, suggesting that GstD3 acts downstream of liposome intracellular processing. As a member of a large family of glutathione S-transferases involved in cellular detoxification and insecticide resistance, GstD3 may have roles in clodronate metabolism that ultimately contribute to its ability to promote apoptosis and cell ablation.

A key step of clodronate toxicity is its incorporation into AMP molecules to form a non-hydrolysable analog of ATP, the adenosine 5’β-γ-dichloromethylene triphosphate (AppCCl2p), that has been shown to inhibit the mitochondrial translocase and putatively induce apoptosis through mitochondrial depolarization. As a result, it has been proposed that aminoacyl-tRNA synthetases could be responsible for the incorporation of clodronate (a bisphosphonate analog of pyrophosphate, PPi) into AMP molecules by a reverse reaction^47^. However, the reaction in which PPi would be incorporated into ADP to regenerate ATP is theoretically possible, but not favored due to thermodynamic and kinetic constraints. In fact, the energy released from ATP hydrolysis and the rapid degradation of PPi by pyrophosphatases ensure that the reverse process does not occur naturally^87^. As a result, aminoacyl-tRNA synthetases (aaRS) typically catalyze the forward reaction of ATP hydrolysis to charge tRNA with an amino acid, producing AMP and pyrophosphate (PPi)^88^. However, the presence of a non-hydrolysable form of PPi, such as clodronate, could hamper the stoichiometry of the reaction and one or multiple enzymes that have PPi and AMP as byproducts, could potentially perform a reverse reaction that incorporates clodronate into AMP molecules. While we did not find aminoacyl-tRNA synthetases in our screen among the enriched hits, if this class of enzymes are involved, a phenotype might not be observable as a result of the essential nature of the aminoacyl-tRNA synthetase involved or because multiple aminoacyl-tRNA synthetases could be catalyzing this reaction creating redundancy and masking the phenotype from genetic enrichment. However, we did observe two nucleotide cyclases among the enriched hits, ADCY5 (AGAP012805) and Gyc89Db (AGAP004564), implicated respectively in the conversion of ATP/GTP to cAMP/cGMP and releasing PPi in the process, yet are not known to catalyze the reverse reaction. Even though these enzymes are not believed to contribute directly to the conversion of clodronate to toxic AppCCl2p, the knockout of these enzymes may be partially protective because of the decreased levels of cAMP/cGTP and the decreased activation of downstream pathways driving cell toxicity. Both cyclic nucleotides are crucial second messengers regulating diverse cellular functions like cellular immunity, autophagy and apoptosis^89^.

While these results enhance our mechanistic understanding of mosquito essential genes and clodronate liposome function, the candidate genes identified in our CRISPR screens will undoubtedly inform a variety of other biological processes that influence mosquito physiology and immune cell function. Based on comparative data in *Drosophila* and human cell lines, we establish a core set of essential genes that can inform further studies on other important mosquito vectors, such as those of the *Aedes* and *Culex* genus. Moreover, the uptake and processing of clodronate liposomes is likely part of a common biological process, such that the genes identified in our screen should provide additional insight into the general mechanisms of phagocytosis and intracellular processing that may inform aspects of host defense, autophagy, apoptosis, and immune cell maturation. Altogether, the results from these initial genome-wide CRISPR screens provide a foundation for additional studies in mosquito cells and *in vivo*, contributing to our further understanding of mosquito biology and mosquito-borne diseases.

## Methods

### Cell culturing

The *Anopheles coluzzii* “screen-ready” (attP+ Cas9+) cell line Sua-5B-IE8-Act::Cas9-2A-Neo^1^ (CVCL_B3N3, Drosophila Genomics Resource Center, stock # 334) as previously described^24^. The cell line was cultured at 25℃ in Schneider’s medium (Gibco), 1x Penicillin-Streptomycin (Gibco) and 10% heat inactivated fetal bovine serum (Gibco) and 500 μg/ml of geneticin (G-418 sulfate, GoldBio).

### Genome-wide library design, and cloning

sgRNAs targeting the whole genome of *Anopheles gambiae* (AgamP4.12) were selected using CRISPR GuideXpress (https://www.flyrnai.org/tools/fly2mosquito/web/) and following the previously described pipeline^24^. Briefly, all computed sgRNAs were retrieved, and the top seven sgRNAs per gene were selected based on the following criteria: minimal OTE (off-target effect) score; maximum ML (machine learning efficiency) score; and filtered to remove sgRNAs that match regions with SNPs in the *Anopheles coluzzii* Sua-5B cell line genome sequence. In addition,sgRNA designs with the BbsI site sequence were removed because BbsI is used for ligation-based cloning into the library vector. The library includes 89,724 unique gene-targeting sgRNAs as well as control and other sgRNAs, as detailed in **Supplementary Table 1**. The sgRNA sequences were cloned into BbsI-digested pLib6.4B-Agam_695 (Accession # OL312683; Addgene # 176668) using the CloneEZ service (Genscript). Cloned vector was subsequently reamplified with a theoretical coverage of <100 times in E. cloni 10GF’ ELITE Electrocompetent Cells (Lucigen) and grown in 500 mL of LB-Ampicillin media at 30°C overnight and bacterial pellets were frozen at -80°C. Before transfection, plasmid DNA was prepared from 50 mL pellets by midiprep (Zymo). Sequencing of the cloned plasmid library confirmed the successful cloning of >98,3% (88159/89711) of designed sgRNAs, detectable with at least one read/sgRNA (circa 94% of guides are detected with at least 10 reads/guide and about 1.7% were lost stochastically).

### Gene essentiality screen

Sua-5B-IE8-Act::Cas9-2A-Neo cells in the log phase of growth were seeded at 35 x 10^6^ cells per 100 mm dish in growth media containing antibiotics. They were transfected with a plasmid mixture containing equimolar amounts of HSP70-ΦC31-Integrase plasmid (pBS130) and sgRNA donor plasmid library (pLib6.4B-Agam_695) using Effectene (Qiagen) according to the manufacturer’s base protocol (“1:25”). We achieved a coverage of ∼ 244 cells/sgRNA by transfecting 735 x 10^6^ cells [90208 sgRNAs x 244 cells/sgRNA x 0.03 (RMCE efficiency) = 735 x 10^6^] in 21 100-mm dishes. After 4 days, each dish was expanded into 2 x 150 cm dishes containing 5 mg/mL puromycin. Cells were cultured for an additional 26 days with media changes and re-seeding every 4 days. Re-seeding at each passage was maintained at a density above 1000 cells/sgRNA to ensure representation of KO pool diversity. Cells were cultured up to 60 days (8 weeks) after transfection. Following selection, genomic DNA was extracted from cell pellets containing >1000 cells/sgRNA using the Quick-gDNA MaxiPrep kit (Zymo). Next, the genomic DNA was barcoded and Illumina sequencing adapters were added via 2-step PCR amplification. Amplicon sequencing was performed using a NextSeq500 at the Biopolymers Facility at Harvard Medical School. Demultiplexing and trimming of barcode labeling was performed using in-house scripts. sgRNAs with a low-read count (<10 reads in the plasmid library) were removed from the readcount files. For identification of base fitness genes the plasmid library vector readcounts from cells after 60 days post-transfection were analyzed with MAGeCK MLE (version 0.5.6) to infer MLE Z-scores for each gene.

To assess the significance of Z-score assignments in inferring true gene essentiality, Z - score average from each replicate was calculated for each gene and plotted against RNAseq expression values obtained from the Sua-5B cell line previously calculated^90^. False-discovery rate (FDR) was inferred from relationships between Z-score and gene expression, as true fitness genes should be among the expressed genes, whereas the identification of a fitness gene that is not expressed represents a false-discovery event. Genes were binned every 5 genes, and the cumulative increase in false-discovery was plotted as a function of Z-score to obtain the FDR. FDR ranking of essential genes revealed 1280 essential genes with 95% confidence. Distribution by Gene Ontology terms of major eukaryotic essential complex components for *Anopheles* within whole genome Z-score distribution was displayed in **Figure 1d**.

### Ortholog mapping and comparison with essential genes in *Drosophila*

Mapping of *Anopheles* genes to *Drosophila* and to human ortholog was done using DIOPT (v 9.0). Ortholog mapping was filtered based on DIOPT rank (only high or moderate rank excluding low rank mapping) and the orthologs of the essential genes in Anopheles were compared with the corresponding data from *Drosophila* or human respectively. Comparisons with a comparable data set from a *Drosophila* CRISPR cell screen were based on MLE Z values from a previous CRISPR screen in S2R+ cells (at the same 5% FDR)^21^. Comparisons with human data were performed using the ‘core essential’ genes identified from human cell lines^46^. The essential genes in *Anopheles* (**Supplementary Table 2**) are compared with *Drosophila* and human essential gene lists in **Supplementary Table 3**.

### Gene set enrichment analysis

To perform gene set enrichment analysis (GSEA), *Drosophila* orthologs mapped as described above from mosquito genes that scored as essential (1280 genes) or ranked within the first fifty hits in the two clodronate liposome screens (88 genes from Clodronate A & B) were used as input for analysis with PANGEA^39^. For essential gene orthologs, gene set enrichment analysis was based on generic gene ontology (GO) slim biological process (BP) terms^40^; Gene List Annotation for Drosophila (GLAD) gene groups^41^, or FlyBase phenotype annotations for classical mutations^42^, and the full sets of outputted enrichment data from PANGEA are included in **Supplementary Table 2**. For clodronate liposome screen hit analysis, the same three gene sets were used, and these were supplemented by additional analysis using the *Drosophila* GO BP and FlyBase Gene Group gene sets (**Supplementary Table 2**). The specific selections made at the PANGEA user interface are indicated on the first row of the PANGEA analysis sheets within **Supplementary Table 2** and **Supplementary Table 5**.

### Positive selection CRISPR screening with clodronate liposomes

For the positive selection screen, 30 days post library transfection cells were selected in media containing puromycin and 16 μM liposome as a control or 8 μM clodrosome. The concentrations used in the screen were established for Sua-5B-IE8-Act::Cas9-2A-Neo cells to be close to the IC50 for the clodrosome (IC50_Clodrosome_=7.4 μm) and negligible for the liposome vehicle (IC50_Liposome_=81.6 μm), as established by assaying total ATP levels (indirect readout of cell grown) during a 6-day treatment using the Cell Titer Glo assay (Promega), as depicted in **Fig. 2A**. Cells were selected through three cycles of treatment. Each cycle of treatment consisted of seeding the cells in media with liposome vehicle or clodronate liposome, followed by media change and re-seeding two additional times. Except in the case of the treatment “Clodronate B,” in which the cells were exposed to selective media a single time for the first 4 days and then allowed to recover with normal media before the next cycle, all the other treatments were performed by continuous exposure to the selective media. The cells were re-seeded at a density above 1000 cells/sgRNA at each passage to ensure representation of KO pool diversity. Following selection, genomic DNA extraction, barcoding, sequencing and analysis was performed as detailed above. Readcount and data analysis, including enrichment analysis and Robust Rank Aggregation score calculation, were performed using MaGeCK 0.5.7 and scatter plots were visualized with Prism (v 10.1.0).

### Mosquito rearing

*Anopheles gambiae* mosquitoes (Keele strain)^91^ were reared at 27°C and 80% relative humidity, with a 14:10 hr light: dark photoperiod. Larvae were fed on commercialized fish flakes (Tetra), while adults were maintained on a 10% sucrose solution and fed on commercial sheep blood (Hemostat) for egg production.

### RNA isolation and gene expression analyses

RNA isolation from whole adult mosquitoes was performed using TRIzol (Invitrogen, Carlsland, CA) according to the manufacturer’s protocol. Two micrograms of total RNA were used for first-strand synthesis using the LunaScript RT SuperMix Kit (NEB). Gene expression was analyzed with quantitative real-time PCR (qPCR) using PowerUp SYBRGreen Master Mix (Thermo Fisher Scientific), while results were analyzed using the 2^-ΔCt^ method and normalized against the internal reference, *rpS7*, as previously described^30,32,92^. All qPCR primers are listed in **Supplementary Table 6**.

### Timing experiments examining the uptake of fluorescent liposome particles

To determine the approximate timing of liposome uptake *in vivo*, mosquitoes were injected with 69 nl of Fluoroliposome-DiO (LP-DiO, Encapsula Nano Sciences) using a 1:50 dilution in 1X PBS. After injection, mosquitoes were incubated at 27°C for 1, 2, 6, 8, or 12 hours, then were injected with a suspension containing 200 μM of Vibrant CM-DiI (Thermo Fisher Scientific) and 2 mM of Hoechst 33342 (Thermo Fisher Scientific) to label mosquito hemocytes. After an additional incubation of 30 min at 27°C to enable *in vivo* staining, hemolymph was perfused from each mosquito using an anticoagulant buffer of 60% v/v Schneider’s Insect medium, 10% v/v fetal bovine serum (FBS), and 30% v/v citrate buffer (98 mM NaOH, 186 mM NaCl, 1.7 mM EDTA, and 41 mM citric acid; buffer pH 4.5) as previously described^30–33,51^. Hemolymph perfusions were placed directly on multi-well microscopic slides for downstream analysis by microscopy. Cells were allowed to adhere for 20 min and fixed with 4% paraformaldehyde (PFA) for 10 min, followed by five washing steps with 1X PBS. Samples were observed under a Zeiss fluorescent microscope to calculate the percentage of hemocytes (of total) taking up the fluorescent LP-DiO particles.

### Phagocyte depletion with clodronate liposomes

Naïve mosquitoes were injected with either clodronate (CLD) or control (LP) liposomes (Encapsula Nano Sciences) to deplete phagocytic immune cell populations in *Anopheles gambiae* as previously described^32,33,51^. Based on the demonstrated IC_50_ of clodronate liposomes *in vitro* as part of this study, liposomes were diluted to a similar concentration using a 1:50 dilution in 1X PBS for all *in vivo* studies herein. Previous studies were performed using a more concentrated 1:5 dilution^32,33,51^. Final concentrations of clodronate liposomes were calculated based on a hemolymph volume of ∼2 μl^93^. To determine the approximate time needed for phagocyte depletion, mosquitoes were injected with either 69nl of CLD or LP, and then incubated at 27°C for 6, 8, 12, or 24 hours. Whole mosquito samples were then processed for RNA isolation and cDNA synthesis as described above. The expression levels of *Eater* and *Nimrod B2* were used as a proxy to demonstrate phagocyte (granulocyte) depletion^32,33,51^.

### dsRNA synthesis and gene-silencing

Candidate genes identified in the genome-wide CRISPR screen were validated using RNAi-mediated gene silencing to confirm their functional roles in the mode of clodronate action. T7 primers specific to each gene (**Supplementary Table 6**) were used to amplify DNA templates from whole female mosquito cDNA samples to synthesize long dsRNAs using the MEGAscript RNAi kit (Thermo Fisher Scientific). Following synthesis, the concentration of dsRNAs was adjusted to 3 μg/μl. Adult female mosquitoes (3-5 days old) were cold anesthetized and injected with 69 nl of dsRNA targeting each candidate gene. For each experiment, mosquitoes were also injected with dsRNA targeting GFP as a negative control. All injections were performed using Nanoject III (Drummond Scientific). Gene-silencing efficiency was evaluated by qPCR 2 days post-injection. All experiments were performed in triplicate.

### Hemolymph perfusion and hemocyte counting

Hemolymph was perfused in adult female *An. gambiae* through the intrathoracic injection of an anticoagulant solution and collection of the perfusate through a small incision in the abdomen as previously described^30,31^. To determine total hemocyte numbers, the collected perfusion from an individual mosquito was added to a disposable Neubauer Improved hemocytometer slide (iNCYTO C-Chip DHC-N01) as previously^29,30,94^.

### Malaria parasite infection

Infections with the rodent malaria model, *Plasmodium berghei*, were performed by first infecting Swiss Webster mice (Charles River) with *P. berghei*-mCherry^95^ parasites as previously described^30,92^. Mosquito infections were performed by allowing mosquitoes to feed on anesthetized *P. berghei*-infected mouse. Following feeding, fully engorged mosquitoes were selected by cold-sorting, then were placed at 19°C. Oocyst numbers were evaluated by fluorescence microscopy in dissected midguts at 10 days post-infection.

### Use of inhibitors to examine liposome uptake

To examine the mechanisms of liposome uptake by mosquito hemocytes, mosquitoes were treated with 200 μM Cytochalasin D (CytoD, Sigma) to inhibit phagocytosis^55–58^ or 25 μg/ml Chlorpromazine hydrochloride (CPZ, MP Biomedical) to impair clathrin-mediated endocytosis^52–54^. Mosquitoes injected with 10% DMSO in 1X PBS were used as negative controls. At 6h post-injection, mosquitoes were injected with a 1:50 dilution of Fluoroliposome-DiO (LP-DiO) in 1X PBS, and then incubated for 8h at 27°C. Following injection with 2 mM Hoechst 33342 to counterstain nuclei, hemolymph was perfused from individual mosquito samples and then observed using a fluorescent microscope to determine the proportions of hemocytes containing fluorescent liposome particles.

Additional experiments were performed to confirm the effects of CytoD on clodronate liposome uptake. Mosquitoes were first injected with 200 μM CytoD or 10% DMSO in 1X PBS and allowed to recover for 6h at 27°C, then followed by injection with control or clodronate liposomes (diluted at 1:50) and incubated at 27°C for 8 hours. The influence of CytoD on clodronate liposome function and resulting phagocyte depletion was evaluated by proxy through the analysis of *Nimrod B2* expression via qPCR^32,33,51^.

### Phagocytosis assays

The effects of candidate genes or inhibitors on phagocytosis were evaluated by injecting adult female mosquitoes with 69 nl of 2% of green fluorescent FluoSpheres (1 μm; Thermo Fisher Scientific) similar to previous studies^30,32,96^. In addition to the beads, mosquitoes were concurrently injected with 100 μM Vibrant CM-DiI and 2 mM of Hoechst 33342 in 1X PBS to counterstain hemocytes, then allowed to recover for 30 min at 27°C. The effects of gene-silencing on phagocytosis were examined approximately 48h after injection with dsRNAs, while the effects of the inhibitor Cytochalasin D were analyzed at 6h post-injection to serve as a positive control to impair phagocytosis^57^. For each experiment, hemolymph was perfused from individual mosquitoes using an anticoagulant buffer and placed on multi-well microscope slides. Hemocytes were allowed to adhere for 20 min and fixed with 4% PFA. Following five washing steps, samples were mounted with Aqua Poly/Mount (Polysciences) and observed under a fluorescent microscope to determine the percentage of phagocytic cells. Approximately 50 hemocytes were counted per individual mosquito, with data collected from two or more replicates (n=16^+^ mosquito samples).

### Use of gene-silencing to examine liposome uptake and processing

To determine the roles of candidate genes on liposome uptake and processing, candidate genes were first silenced by the injection of dsRNA in naive adult female mosquitoes. Two days post-injection, gene-silenced mosquitoes were injected with a 1:50 dilution of Fluoroliposome-DiO in 1X PBS. Following incubation for 8 hours, phenotypes were evaluated in individual mosquitoes as the percentage of hemocytes containing liposome particles (LP-DiO^+^) to evaluate liposome uptake, or as diffused patterns of DiO (DiO^+^) in the cytosol that support liposome processing and degradation.

### Immunofluorescence of cellular localization

To visualize the co-localization of liposome particles with the lysosome, mosquitoes were perfused with an anticoagulant buffer at 8h post-injection with a 1:50 dilution of LP-DiO. Hemocytes were allowed to adhere for 30 min without fixation, and then incubated with the lysosome-specific dye, LysoView 594 (Biotium), using a 1:500 dilution in 1X PBS for 1 hour. Samples were mounted with ProLongDiamond AntiFade Mountant with DAPI (Life Technologies) and immediately observed using fluorescence microscopy (Zeiss Axio Imager.M2).

### Use of inhibitors to impair lysosome acidification

To investigate the involvement of lysosome function in regulating the processing of clodronate-liposomes, mosquitoes were treated with 25 μM of Bafilomycin A1 (BafA1; Cayman), a proton pump V-ATPase inhibitor, or 10% DMSO in 1X PBS. Mosquitoes were incubated for 16h at 27°C as previously described^97^, then injected with Fluoroliposome-DiO using a 1:50 dilution in 1X PBS. The number of hemocytes displaying intact liposome particles (LP-DiO^+^) or diffused patterns of DiO (DiO^+^) in the cytosol was determined by immunofluorescence. To examine the effects of BafA1 on clodronate function, mosquitoes were injected with clodronate or control liposomes at 1:50 dilution in 1X PBS following treatment with Baf A1. At 8h post-injection, phagocyte depletion was evaluated by proxy through *Nimrod B2* expression via qPCR^32,33,51^.

## Supporting information

Supporting Information containing Supplementary Figures 1-6

## Acknowledgments

We would like to thank Ian Schneider for kindly providing the cytochalasin D inhibitor. Work at the Drosophila Research & Screening Center-Biomedical Technology Research Resource (DRSC-BTRR) was supported by NIH NIGMS P41 GM132087 to NP and SEM. Additional support was provided by NIH R21 AI166857 to RCS, while DRH is supported by the National Science Foundation Graduate Research Fellowship Program under Grant No. 2336877. NP is an Investigator of the Howard Hughes Medical Institute. This article is subject to HHMI’s Open Access to Publications policy. HHMI lab heads have previously granted a nonexclusive CC BY 4.0 license to the public and a sublicensable license to HHMI in their research articles. Pursuant to those licenses, the author-accepted manuscript of this article can be made freely available under a CC BY 4.0 license immediately upon publication.

## Notes

### Competing Interest Statement

The authors have declared no competing interest.

