## Supporting Information containing Supplementary Figures 1-6 for "A genome-wide CRISPR screen in *Anopheles* mosquito cells identifies essential genes and required components of clodronate liposome function"

### Included supporting information

#### *Supplementary Figures*

**Supplementary Fig. 1.** Gene set enrichment analysis of *Drosophila* orthologs of *Anopheles* essential genes using the PANGEA online resource.

**Supplementary Fig. 2**. *Serpent*-silencing influences mosquito immune cell numbers and malaria parasite infection.

**Supplementary Fig. 3.** Gene set enrichment analysis of *Drosophila* orthologs of *Anopheles* clodronate resistance screen results using the PANGEA online resource.

**Supplementary Fig. 4.** Phagocyte depletion using different concentrations of clodronate liposomes.

**Supplementary Fig. 5.** Timing of liposome uptake and immune cell depletion.

**Supplementary Fig. 6.** Day 4 RNAi of candidate genes.

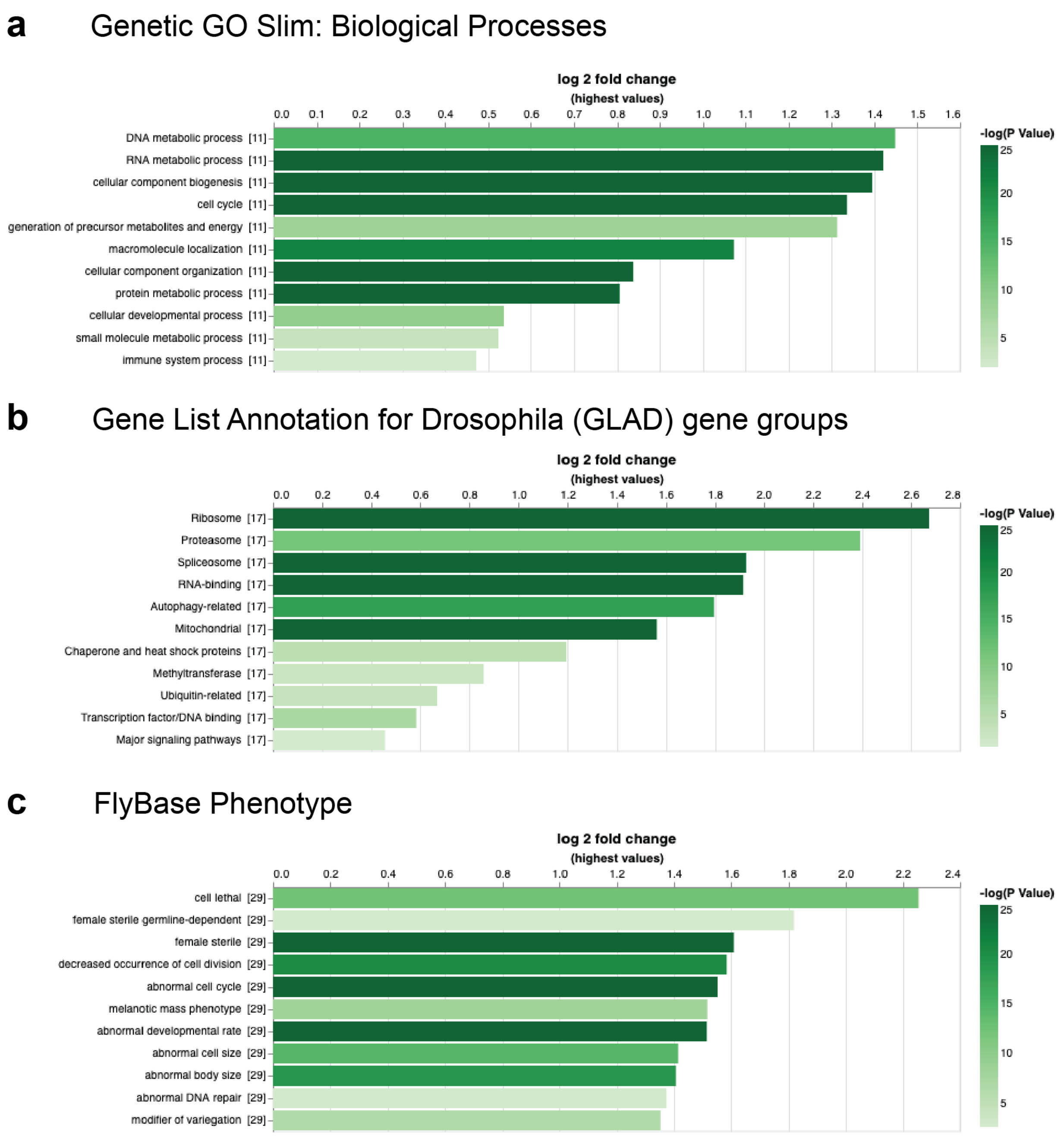


**Supplementary Fig. 1. Gene set enrichment analysis of *Drosophila* orthologs of *Anopheles* essential genes using the PANGEA online resource.** Genes that scored as fitness genes/essential genes in the genome-wide *Anopheles* cell screen were mapped to *Drosophila* orthologs using DIOPT. Only the top-scoring ortholog match was selected; if more than one *Drosophila* gene had the same DIOPT ortholog score, we arbitrarily chose one of the genes for the analysis. The *Drosophila* gene list was then used as the input at PANGEA. **(a)** Enrichment of gene ontology (GO) slim "Biological Process" annotations (Gene Ontology sets>Gene Ontology Subsets>Generic GO consortium Subsets (GO slim)>SLIM1 GO BP). **(b)** Enrichment of Gene List Annotation for Drosophila (GLAD) gene sets (Other Gene sets>DRSC GLAD Gene Group). **(c)** Enrichment of FlyBase phenotype annotations for classical mutations (Other Gene sets>Phenotype>FlyBase phenotype for classical alleles). In each case, the top ten results as ranked based on fold-change are displayed. Length of the bars represents fold enrichment of genes within each category while darkness reflects the *P* value. Only GO terms with *P* value below the 0.05 threshold are displayed, up to a maximum of 10 terms. The full PANGEA analysis outputs including results of multiple statistical analyses are included in **Supplementary Table 2**.

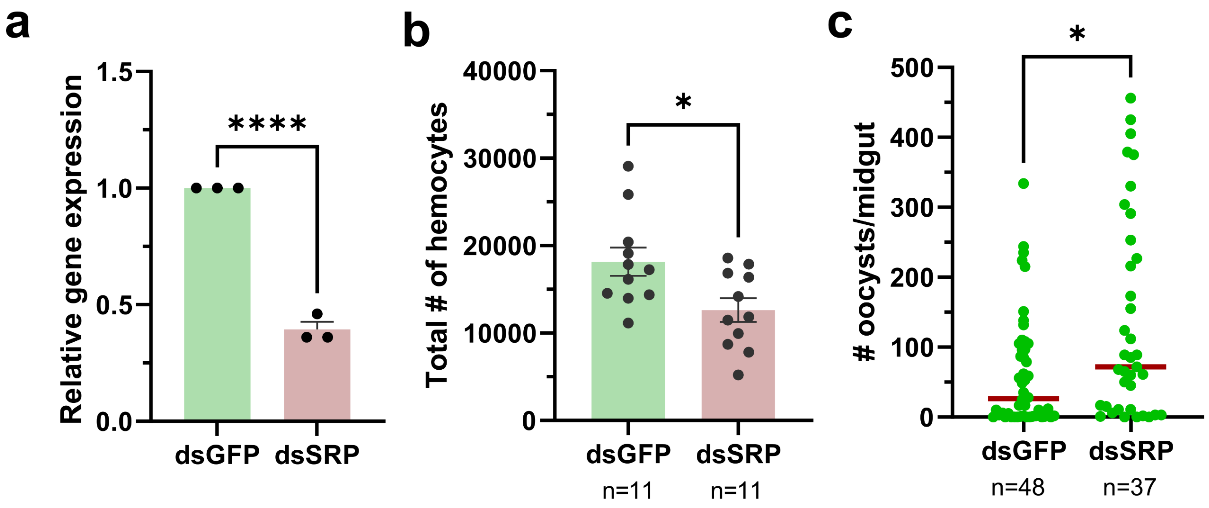


**Supplementary Fig. 2**. ***Serpent*-silencing influences mosquito immune cell numbers and malaria parasite infection.** (**a**) Confirmation of *serpent* (*srp*) gene-silencing by qRT-PCR. Data were collected from three independent experiments of ~10 adult female mosquitoes at two days post-dsRNA injection. Statistical analysis was performed using an unpaired t-test. (**b**) Examinations of total hemocyte counts from individual mosquitoes (n=11) were performed in control (*dsGFP*) and *srp*-silenced backgrounds. (**c**) The effects of *srp*-silencing on *P. berghei* infection were evaluated by oocyst numbers at 10 days post-infection. Data in **b** and **c** were analyzed by Mann-Whitney to determine significance. For all experiments, significant differences are indicated by asterisks (*, $P < 0.05$; ****, $P < 0.0001$).

GO: Biological Processes

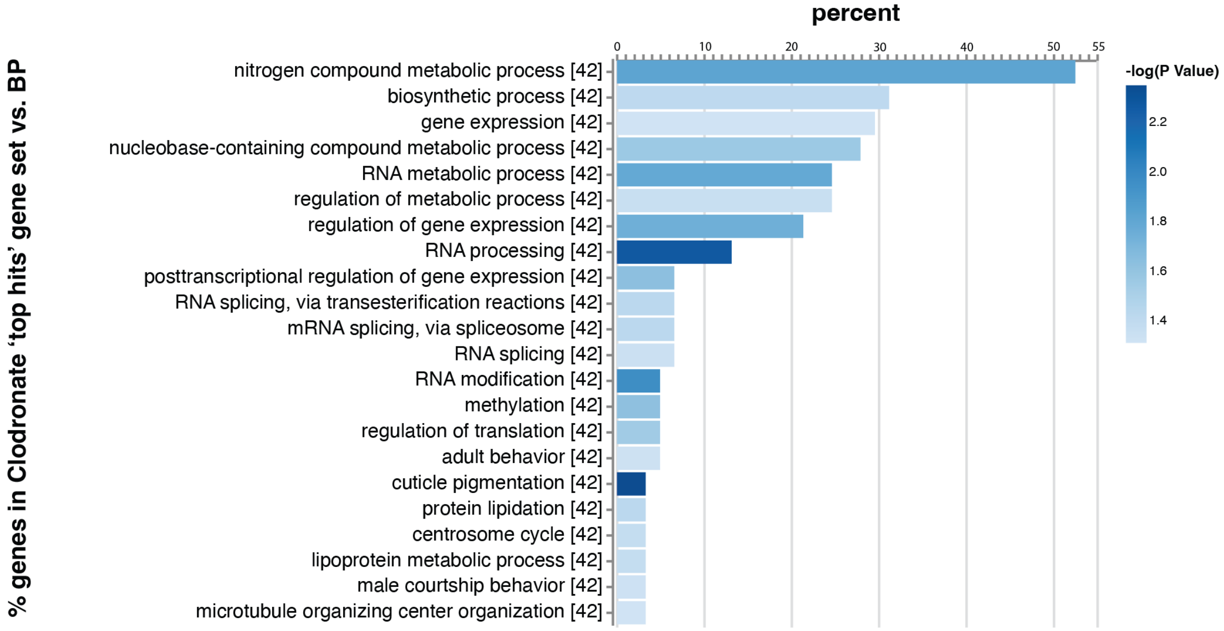


**Supplementary Fig. 3. Gene set enrichment analysis of *Drosophila* orthologs of *Anopheles* clodronate resistance screen results using the PANGEA online resource.** Genes that scored in the genome-wide *Anopheles* cell screen for resistance to clodronate treatment were mapped to *Drosophila* orthologs using DIOPT. Only the top-scoring ortholog match was selected; if more than one *Drosophila* gene had the same DIOPT ortholog score, we arbitrarily chose one of the genes for the analysis. The *Drosophila* gene list was then used as the input at PANGEA. Only significantly enriched gene sets ($p<0.05$) are shown for enrichment of GO terms for *Drosophila* as reported by FlyBase and the Alliance of Genome Resources (Gene Ontology sets> GO Biological Processes). The enriched gene sets are ranked based on the percentage of genes in the clodronate screen gene set Vs the GO Biological Processes gene set. Length of the bars represents percentage of genes within each category while darkness reflects the *P* value calculated on the GO enrichment. The full PANGEA analysis outputs including results of multiple statistical analyses are included in **Supplementary Table 5**.

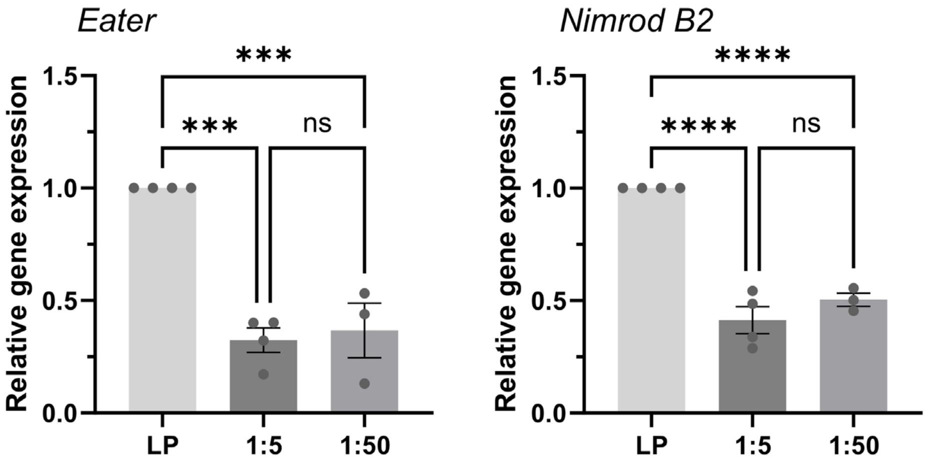


**Supplementary Fig. 4. Phagocyte depletion using different concentrations of clodronate liposomes.** The expression of *eater* and *Nimrod B2* was evaluated by qRT-PCR in mosquitoes injected with 1:5 or 1:50 dilutions of clodronate liposomes and compared to empty liposome controls (LP). Pooled mosquitoes from three or more independent experiments were examined at 24 hours post-injection using a one-way ANOVA and a Holm-Sidak's multiple comparison test. Significant differences are indicated by asterisks (***, $P < 0.001$; ****, $P < 0.0001$). ns, not significant.

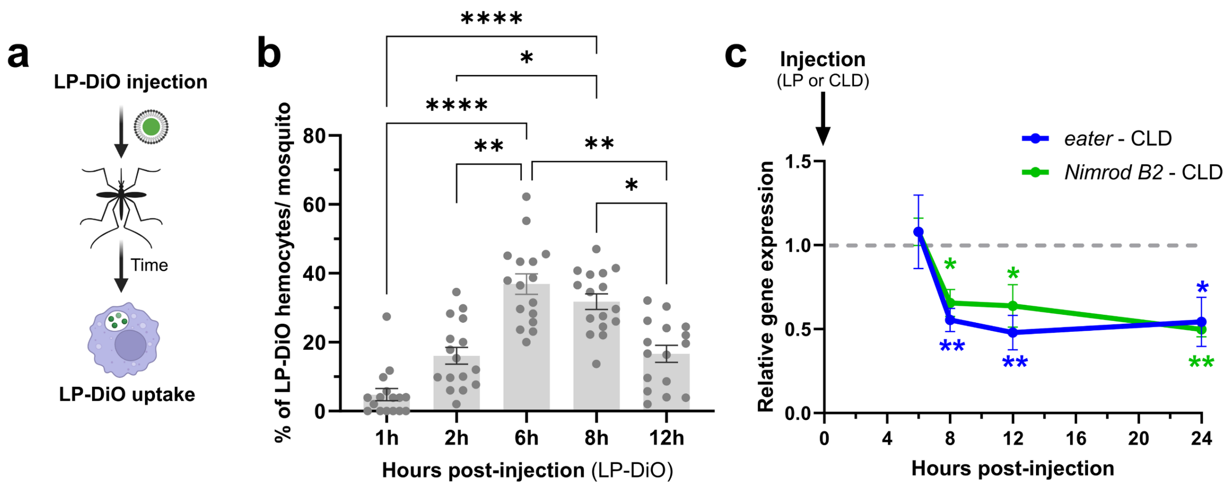


**Supplementary Fig. 5. Timing of liposome uptake and immune cell depletion.** (**a**) Experimental overview of methodology used to examine the timing of liposome (LP-DiO) uptake in vivo by circulating immune cells. **(b)** The percentage of LP-DiO$^+$ cells at 1, 2, 6, 8, or 12 hours post-injection. Each dot corresponds to data collected from an individual mosquito. Data were analyzed using Kruskal-Wallis and Dunn's multiple comparison test to determine significance. **(c)** The timing of clodronate liposome-mediated cell ablation was determined *in vivo* by measuring *eater* and *Nimrod B2* expression at 6, 8, 12, and 24 hours post-injection of control or clodronate liposomes in adult female mosquitoes. Data from three or more independent experiments were analyzed using an unpaired t test to compare expression levels between control- or clodronate liposome-treated mosquitoes. Significant differences are indicated by asterisks (*, $P$ < 0.05; **, $P$ < 0.01; ****, $P$ < 0.0001).

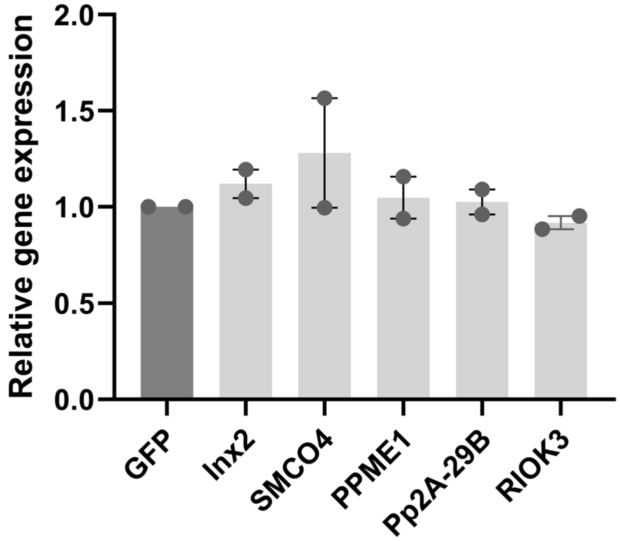


**Supplementary Fig. 6. Day 4 RNAi of candidate genes.** For candidate genes where RNAi was unsuccessful at two days post-injection, additional RNAi experiments were performed in which knockdowns were evaluated at four days post-injection. Expression data from two independent experiments are displayed as the mean ±SEM and compared to GFP controls.
