## Supplementary Table 1 for "A genome-wide CRISPR screen in *Anopheles* mosquito cells identifies essential genes and required components of clodronate liposome function"

**Supplementary Table 1.** Makeup of the genome-wide sgRNA library used in this study.

| **Guide type or target** | **Number of guides (number of genes)** |
| --- | --- |
| Targeting intergenic regions (negative controls) | 400 sgRNAs |
| Non-targeting (negative controls) | 100 sgRNAs |
| Positive controls (based on Viswanatha, Mameli, et al. 2021 *Nature Communications*) | 461 sgRNAs (3 genes) |
| Protein-coding genes (total unique) | 87,812 sgRNAs (12,774 genes)   - 12,323 genes covered by 7 sgRNAs/gene - 301 genes covered by 3-6 sgRNAs/gene - 150 genes covered by 1-2 sgRNAs/gene |
| Non-protein-coding genes (total unique) | 951 sgRNAs (230 genes)   - 57 genes covered by 7 sgRNAs/gene - 104 genes covered by 3-6 sgRNAs/gene - 69 genes covered by 1-2 sgRNAs/gene |
| Duplicated designs* | 484 sgRNAs (235 genes) |
| Total unique designs | 89,724 sgRNAs |
| Total sgRNAs including duplications | 90,208 sgRNAs |

* We included two copies of one sgRNA design for 71 genes for which only 1 unique sgRNA per gene met our design criteria, 79 genes for which only 2 sgRNAs met criteria, and 85 genes for which only 3 sgRNAs met criteria.
